# Live Virus Neutralisation of the 501Y.V1 and 501Y.V2 SARS-CoV-2 Variants following INO-4800 Vaccination of Ferrets

**DOI:** 10.1101/2021.04.17.440246

**Authors:** Shane Riddell, Sarah Goldie, Alexander J. McAuley, Michael J. Kuiper, Peter A. Durr, Kim Blasdell, Mary Tachedjian, Julian D. Druce, Trevor R.F. Smith, Kate E. Broderick, Seshadri S. Vasan

## Abstract

The ongoing COVID-19 pandemic has resulted in significant global morbidity and mortality on a scale similar to the influenza pandemic of 1918. Over the course of the last few months, a number of SARS-CoV-2 variants have been identified against which vaccine-induced immune responses may be less effective. These “variants-of-concern” have garnered significant attention in the media, with discussion around their impact on the future of the pandemic and the ability of leading COVID-19 vaccines to protect against them effectively. To address concerns about emerging SARS-CoV-2 variants affecting vaccine-induced immunity, we investigated the neutralisation of representative ‘G614’, ‘501Y.V1’ and ‘501Y.V2’ virus isolates using sera from ferrets that had received prime-boost doses of the DNA vaccine, INO-4800. Neutralisation titres against G614 and 501Y.V1 were comparable, but titres against the 501Y.V2 variant were approximately 4-fold lower, similar to results reported with other nucleic acid vaccines and supported by *in silico* biomolecular modelling. The results confirm that the vaccine-induced neutralising antibodies generated by INO-4800 remain effective against current variants-of-concern, albeit with lower neutralisation titres against 501Y.V2 similar to other leading nucleic acid-based vaccines.

## Introduction

With the continuing roll-out of vaccines against severe acute respiratory syndrome coronavirus 2 (SARS-CoV-2), primarily developed against the ancestral ‘D614’ version of the virus^1^, there is concern that emerging ‘variants-of-concern’ and ‘variants-of-interest’ may have acquired mutations resulting in lower neutralisation by infection- and vaccine-induced antibodies. Neutralising antibodies are a key correlate of protection against SARS-CoV-2 in humans and non-human primates^2-4^, so the determination of antibody-mediated neutralisation against emerging variants is important when assessing vaccine efficacy. Recent publications have demonstrated that these concerns have some basis, with reduced neutralisation of both the ‘501Y.V1’ variant (also called the ‘UK variant’ or ‘B.1.1.7’) and the ‘501Y.V2’ variant (also called the ‘South African variant’ or ‘B.1.351’) by leading mRNA-based vaccines^5^. Similarly, neutralisation studies using human serum samples from coronavirus disease-19 (COVID-19) patients in South Africa reveal significant decreases in neutralisation efficiency against the 501Y.V2 variant^6,7^.

We recently investigated the neutralisation of SARS-CoV-2 isolates containing either the ‘D614’ or the ‘G614’ variation of the Spike protein, using serum samples from ferrets vaccinated with INO-4800, a leading DNA vaccine candidate against SARS-CoV-2^8,9^, demonstrating the D614G mutation was unlikely to affect vaccine-induced antibody-mediated virus neutralisation. Given the interest around the 501Y.V1 and 501Y.V2 variants, we decided to follow up our previous study by comparing neutralisation titres of INO-4800-vaccinated ferret samples against representative SARS-CoV-2 isolates. We believe such a study is important because: a) DNA vaccines such as INO-4800 can be rapidly modified similar to mRNA vaccines; b) INO-4800 remains stable at room temperature for an extended period, making it especially suitable for combatting the pandemic in resource-poor and remote settings^9^; and, c) INO-4800 has a favourable safety and reactogenicity profile in humans, as demonstrated in clinical trials.

## Materials and Methods

### Viruses, vaccine, and cells

Australian SARS-CoV-2 isolates hCoV-19/Australia/VIC31/2020 (VIC31) and hCoV-19/Australia/VIC17990/2020 (VIC17990; 501Y.V1 variant) were provided as Passage 1 material by the Victorian Infectious Diseases Reference Laboratory (VIDRL), Melbourne, Australia. hCoV-19/South Africa/KRISP-K005325/2020 (501Y.V2.HV001; 501Y.V2 variant) was provided as Passage 3 material from the Africa Health Research Institute, Durban, South Africa. INO-4800 was provided by Inovio Pharmaceuticals, San Diego, USA, for preclinical testing within the ferret model of COVID-19. This DNA vaccine is a plasmid containing a coding sequence for SARS-CoV-2 (Wuhan-Hu-1) Spike glycoprotein without constraints on protein folding^9^.

VIC31 Passage 2, VIC17990 Passage 2, and 501Y.V2.HV001 Passage 4 virus stocks were grown in VeroE6 cells (CCL81; American Type Culture Collection (ATCC), Manassas, VA, USA). Briefly, VeroE6 cells were grown in 150cm^2^ flasks in Dulbecco’s Modified Eagle Medium (DMEM) containing 10% heat-inactivated foetal bovine serum (FBS), 10mM HEPES, 100U/mL penicillin, 100μg/mL streptomycin, and 250ng/mL amphotericin B (all components from ThermoFisher Scientific, Scoresby, VIC, Australia) until 79-90% confluent. The received SARS-CoV-2 isolates were diluted in DMEM containing 10mM HEPES, 100U/mL penicillin, 100μg/mL streptomycin, and 250ng/mL amphotericin B, but no FBS (DMEM-D). Cells were inoculated with 4mL diluted virus and were incubated for 30min at 37°C/5% CO_2_ before 40mL DMEM containing 2% FBS, 10mM HEPES, 100U/mL penicillin, 100μg/mL streptomycin, and 250ng/mL amphotericin B was added. The flasks were incubated for an additional 48h before supernatant was harvested. ATCC VeroE6 cells were additionally used for virus neutralisation assays (see below).

### Next-Generation Sequencing

Virus stocks were sequenced using a MiniSeq platform (Illumina, Inc; San Diego, CA, USA). In brief, 100 µL cell culture supernatant from the infected Vero E6 cells was combined with 300 µL TRIzol reagent (Thermo Fisher Scientific) and RNA was purified using a Direct-zol RNA Miniprep kit (Zymo Research; Irvine, CA, USA). Purified RNA was further concentrated using an RNA Clean-and-Concentrator kit (Zymo Research), followed by quantification on a DeNovix DS-11 FX Fluorometer. RNA was converted to double-stranded cDNA, ligated then isothermally amplified using a QIAseq FX single cell RNA library kit (Qiagen). Fragmentation and dual-index library preparation was conducted with an Illumina DNA Prep, Tagmentation Library Preparation kit. Average library size was determined using a Bioanalyser (Agilent Technologies; San Diego, CA, USA) and quantified with a Qubit 3.0 Fluorometer (Invitrogen; Carlsbad, CA, USA). Denatured libraries were sequenced on an Illumina MiniSeq using a 300-cycle Mid-Output Reagent kit as per the manufacturer’s protocol. Paired-end Fastq reads were trimmed for quality and mapped to the published sequence for the SARS-CoV-2 reference isolate Wuhan-Hu-1 (RefSeq: NC_045512.2) using CLC Genomics Workbench version 21. Variations from the reference sequence were determined with the CLC Genomics Workbench Basic Variant Detection tool set at a minimum frequency of equal to or greater than 35%.

### Serum samples

Serum samples used for neutralisation assays were generated and selected with the consent of the sponsor (Coalition for Epidemic Preparedness Innovations), as previously described^8^. In brief, four male and four female ferrets received two doses of INO-4800 via intramuscular administration of 1 mg plasmid DNA to the caudal thigh muscle, followed by electroporation split across two sites using the CELLECTRA® device. The prime dose was given on day 0 and the boost on day 28. Sera collected on day 35 or day 42 from three male and three female ferrets were chosen at random to allow for sufficient seroconversion, and to ensure that no ferret provided more than one test sample. Two unvaccinated ferret sera were also included as negative control sera, to ensure that there was no non-specific neutralisation of the viruses. Parent vaccine efficacy studies were approved by the Animal Ethics Committee at CSIRO ACDP (Approval Reference: AEC 2004). No additional ethics approval was required to perform neutralisation assays on serum samples collected from the parent study.

### Serum dilution

Each serum sample was diluted 1:20 in DMEM-D (see cell culture methods above) in a deep-well plate on a single occasion, followed by a 2-fold serial dilution in medium across the plate up to 1:20,480. The dilution series for each serum sample was dispensed into triplicate rows of a 96-well plate, for a total volume of 50μL per well and triplicate wells per sample dilution.

### Neutralisation assay

For the serum-containing wells, 50μL virus diluted in medium to contain approximately 100 TCID_50_(checked by back-titration) was added to each well. The plates were incubated at 37°C/5% CO_2_ for 1h to allow neutralisation complexes to form between the antibodies and the virus. At the end of the incubation, 100μL VeroE6 cells (propagated as outlined above for virus stock generation) were added to each well and the plates returned to the incubator for 4 days. Each well was scored for the presence of viral CPE, readily discernible on Day 4 post-infection, with neutralisation titres assigned to each serum replicate based upon the highest dilution that prevented discernible cytopathic effect.

### Statistical analysis

Mixed effects analysis of variance (ANOVA) was used to assess differences between SARS-CoV-2 isolates (the fixed effect), with the random effect being the 3 test replicates. For the *post hoc* analysis to detect significant factor level differences we undertook pairwise comparisons with Tukey’s adjustment. All analyses were undertaken in R 4.0, using the *nlme v. 3*.*1*. package for the mixed effects ANOVA modelling and *multcomp v. 1*.*4* for the *post hoc* comparisons.

### Molecular modelling

Fully glycosylated models of the Spike protein G614, 501Y.V1 and 501Y.V2 variants were built based on ‘6VSB’ protein databank (PDB) structure^10^ minus transmembrane domains (residues 1161-1272) and included one ACE2 protein bound to the receptor-binding domain. All models were simulated in aqueous solution (TIP3 water, 310K, 0.15M ions, NVT ensemble, each approximately 725,000 atoms in size) using the software NAMD 2.14^11^ for 150 nanoseconds each. Models were visualized with VMD and Nanome^12,13^.

## Results

Stocks of VIC31 (hCoV-19/Australia/VIC31/2020; EPI_ISL_419750), the G614 isolate used in our previous study^8^; VIC17990 (hCoV-19/Australia/VIC17990/2020; EPI_ISL_779605), an Australian isolate of the 501Y.V1 variant; and 501Y.V2.HV001 (hCoV-19/South Africa/KRISP-K005325/2020; EPI_ISL_678615), a South African isolate of the 501Y.V2 variant, were propagated and titrated in Vero E6 cells prior to use, with genome sequences confirmed by next-generation sequencing. Mutations within the Spike protein (relative to the reference D614 isolate, Wuhan-Hu-1; RefSeq: NC_045512.2) can be found in **Table 1**.

**Table 1:**
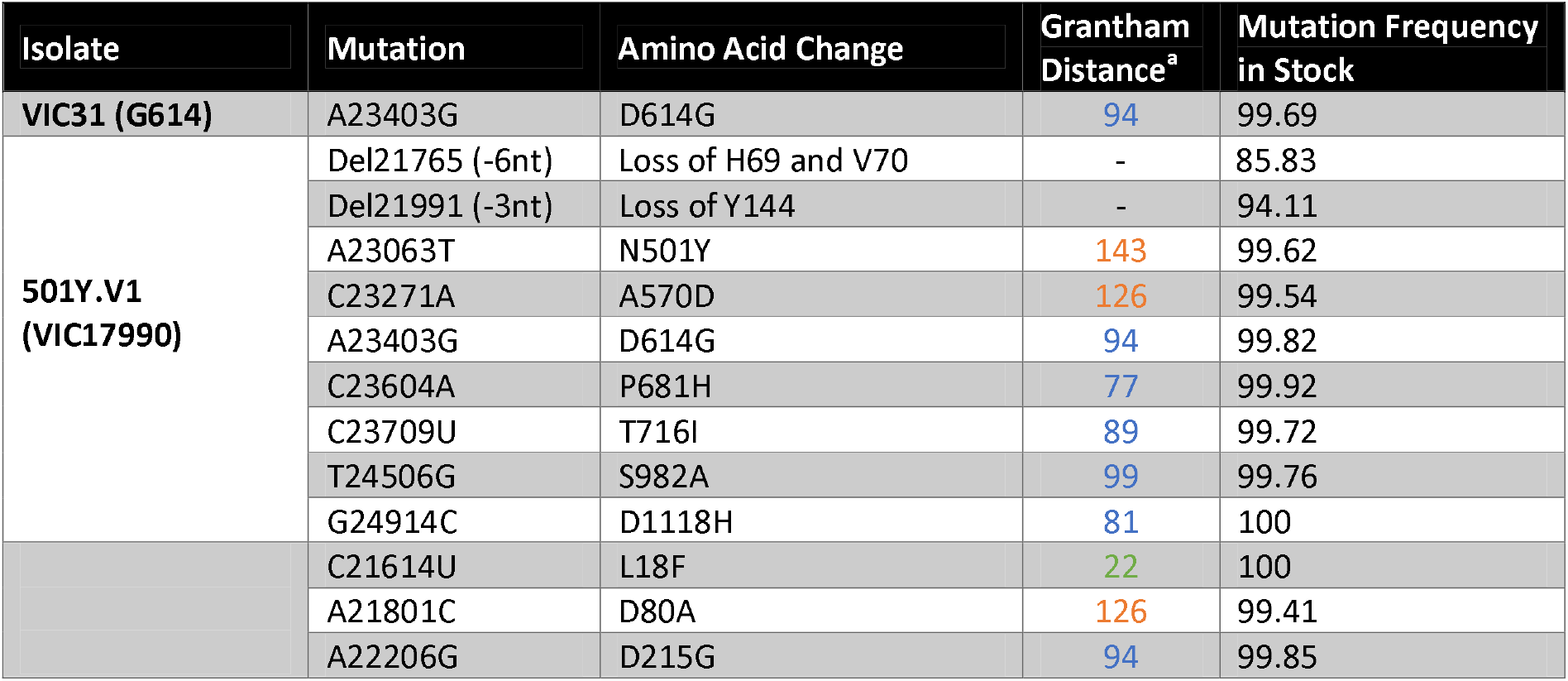
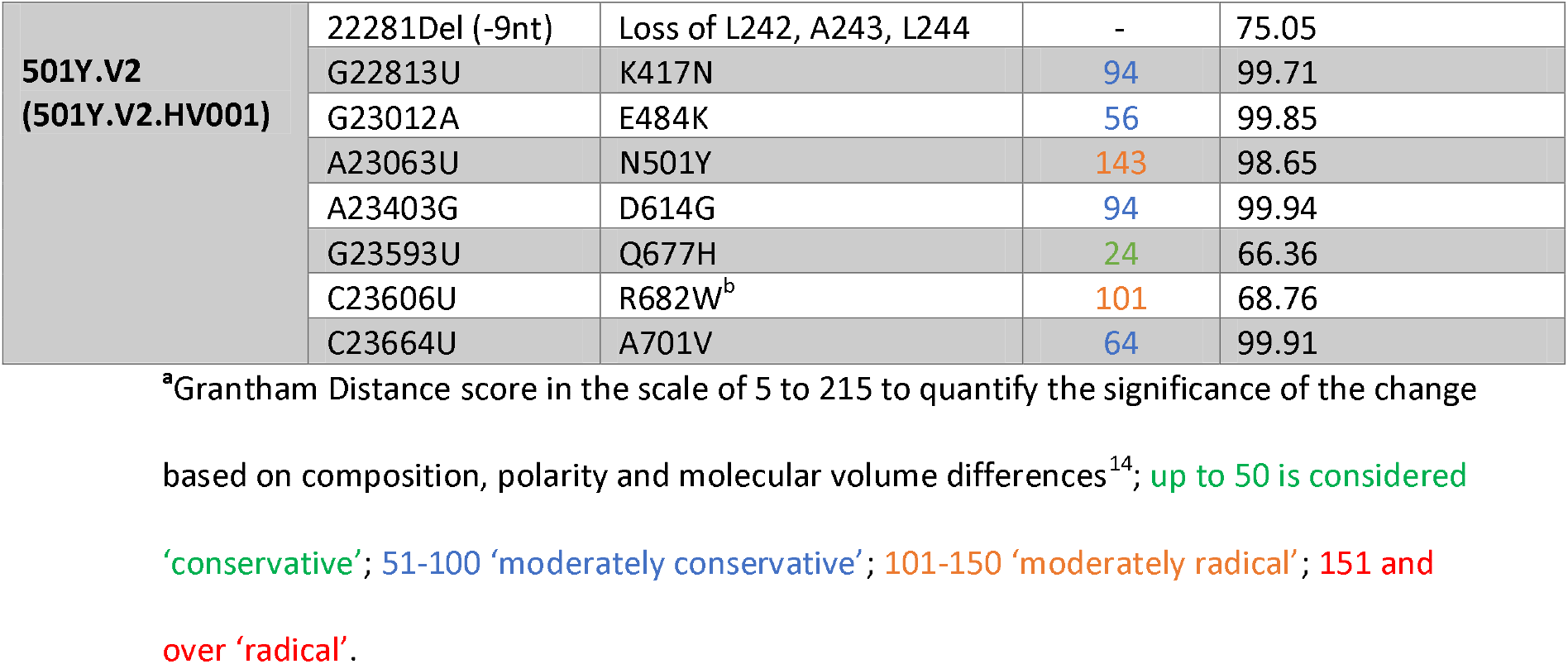
Mutations in Spike Protein Present in Variant Stocks (Compared to the Wuhan-Hu-1 Reference Sequence)

For consistency and comparison with McAuley et al^8^, microneutralisation assays were performed in triplicate for each serum/isolate combination, using the same three male and three female ferret samples used previously. We additionally included negative control (unvaccinated) ferret serum samples. Neutralisation titres against VIC31 were similar between this study and the previous one^8^ (mean neutralisation titre of 90 for previous study; 97 for current study), with no significant difference by mixed effects ANOVA (p>0.05), confirming inter-assay reproducibility **(Figure 1a)**. Mean neutralisation titres were calculated for the different isolates on log_2_-transformed data: 97 for VIC31; 105 for 501Y.V1; 24 for 501Y.V2. Mixed effects ANOVA comparison of the neutralisation titres against the three isolates revealed no significant difference in neutralisation of the 501Y.V2 isolate (VIC17990) compared to VIC31, however titres against 501Y.V2 were significantly lower than against the other two (p<0.001; 25% of the VIC31 value; 23% of the 501Y.V1 value) **(Figure 1b)**.

**Figure 1:**
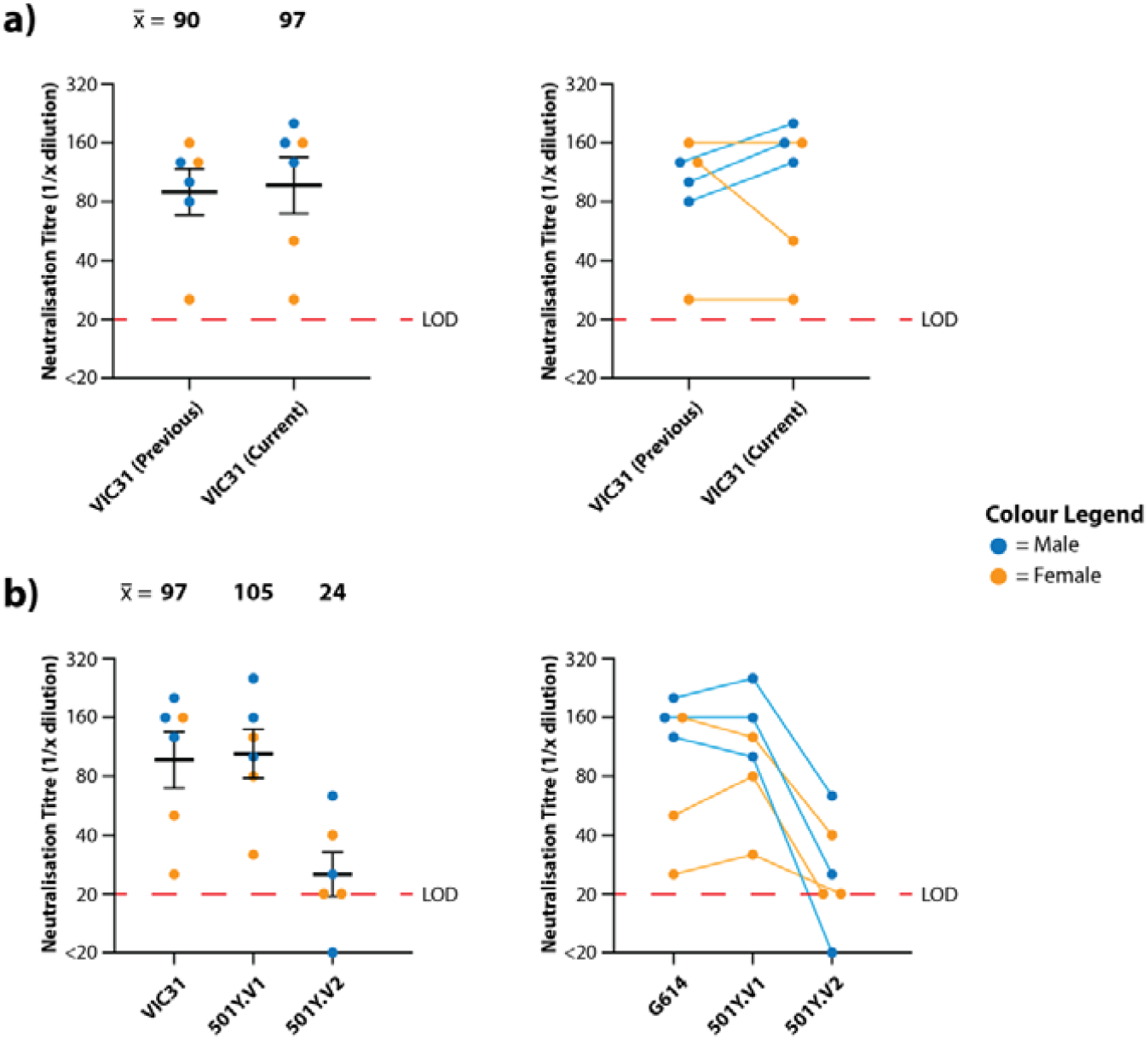
Neutralisation of SARS-CoV-2 Variants-of-Concern with Serum from INO-4800-Vaccinated Ferrets. Neutralisation titres were obtained in triplicate for serum samples collected from ferrets receiving prime and boost doses of INO-4800. a) Comparison of anti-VIC31 (G614 variant) titres from the current and previous studies^8^ demonstrating inter-assay reproducibility (left; p>0.05 by paired, two-tailed t-test), and tracking of values from individual animals (right). b) Comparison of neutralisation titres against VIC31 (G614 variant), 501Y.V1 and 501Y.V2 variants (left; p<0.001 by mixed effects ANOVA), and tracking of values from individual animals (right). Horizonal lines represent geometric mean and geometric standard error of the mean.

Molecular modelling of the variants shows more cumulative structural changes in 501Y.V2 than 501Y.V1 (**Figure 2; Table 1**), consistent with observed lower neutralisation titres. The receptor-binding domain (RBD) of the Spike protein in the 501Y.V1 variant contains the N501Y mutation, which has been shown to increase binding to the ACE2 receptor^15,16^, while 501Y.V2 additionally contains the mutations K417N and E484K in the RBD **(Figure 2b,c)**. The E484K mutation, although a moderately conservative change in Grantham Distance, has been shown to be an extremely potent escape mutant, showing resistance to many monoclonal and even polyclonal antibodies^17^. The K417N mutation, also considered a moderate conservative change, appears to increase hydrogen bonding interactions with the ACE2 receptor via ACE2 N-linked glycosylation at asparagine position Asn90 **(Supplementary Figure S1)**. The N-terminal domain (NTD) of the Spike protein in the 501Y.V1 variant contains two deletions at residue positions 69-70 and 144, potentially modulating the Spike protein’s glycosylation arrangements attached at asparagine residues Asn74 and Asn149. More extensive changes to the NTD exist in the 501Y.V2 variant, with a cluster of mutations (within approximately 15Å radius of D80A), including L18F, the moderately radical D80A, the moderately conservative D215G, and a 242-244 deletion **(Figure 2d,e)**. Key implications of these changes within the NTD are: (a) the D80A and D215G mutations in 501Y.V2 have replaced aspartic acid residues which can stabilize salt bridge interactions with the arginine residues Arg78 and Arg214 respectively **(Supplementary Figure S2)**; and (b) the 242-244 deletion has removed the ‘Leu-Ala-Leu’ residues, causing local residue sidechain rearrangement and shortening a solvent exposed loop (residues 245-259), likely altering its antigenic presentation. These combined structural changes in 501Y.V2 could explain our observed reduced inhibition potency relative to the 501Y.V1 and D614G variants.

**Figure 2:**
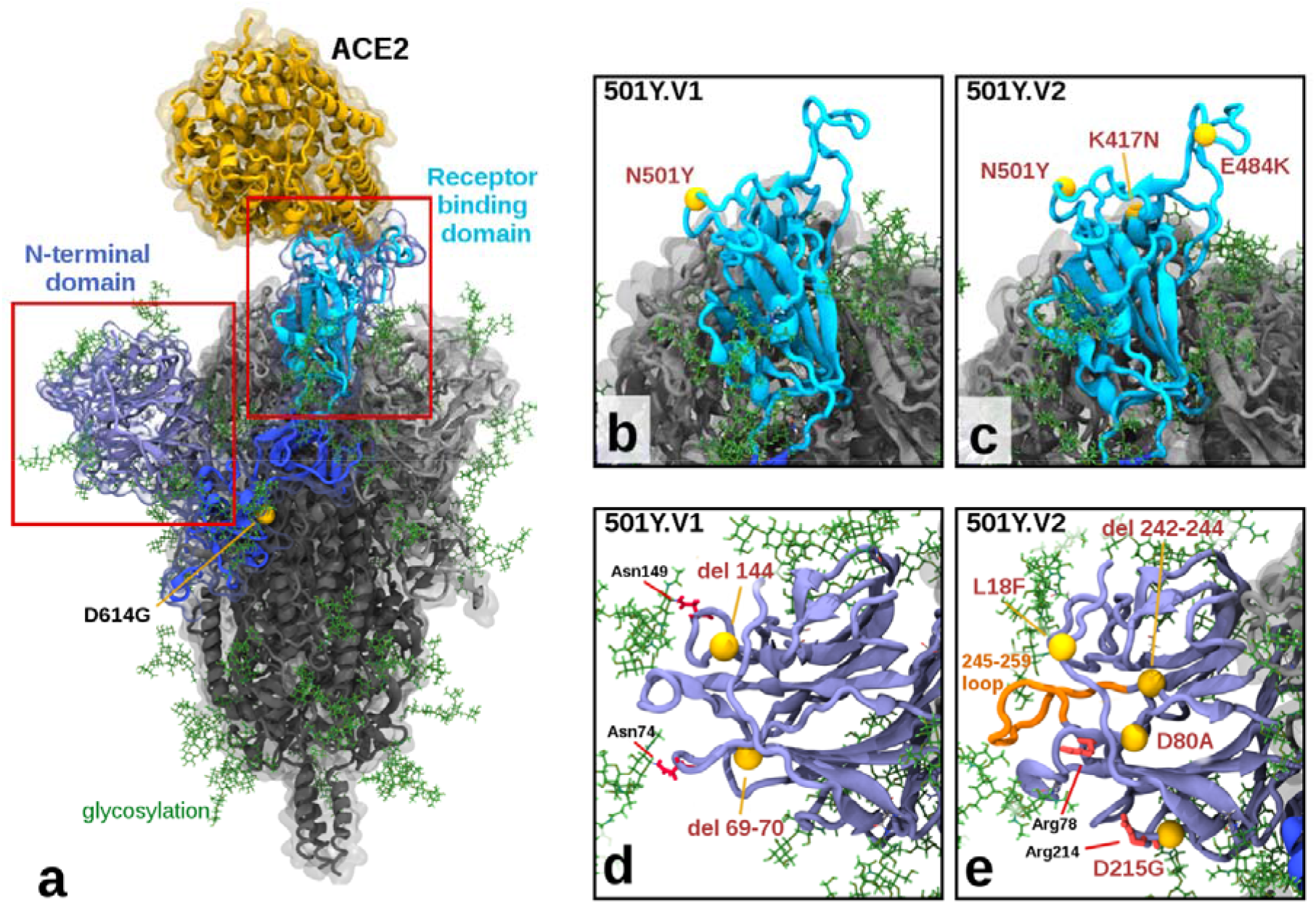
Illustration of Mutations in 501Y.V1 and 501Y.V2 Relative to the Ancestral Form. a) Structure of the glycosylated SARS CoV2 Spike protein highlighting a S1 monomer (in blues) and relative positions of the N terminal domain (NTD) and the receptor-binding domain (RBD) and the bound ACE2 receptor (in yellow). The position of D614G (common to all tested variants) is also highlighted. b) & c) Side by side comparison of 501Y.V1 and 501Y.V2 variants in the RBD, showing the V2 variant having additional K417N and E484K mutations. d) & e) Side by side comparisons of 501Y.V1 and 501Y.V2 variants in the NTD, showing relative locations of mutations and deletions. (Model files available in Supplementary Materials).

## Discussion

The results of this study are consistent with previous analyses of the effect of emerging SARS-CoV-2 variants on infection- and vaccine-induced immune responses ^5-7^. Indeed, the reductions in neutralisation titres (and by extension, vaccine-induced protection) observed with the 501Y.V2 variant are comparable for leading nucleic acid based COVID-19 vaccines – approximately 4-fold for INO-4800 (Inovio Pharmaceuticals), compared to 6.5-fold for BNT162b2 (BioNTech/Pfizer) and 8.6-fold for mRNA-1273 (Moderna) – and substantially less than the 86-fold reduction reported for AZD1222 (Oxford/AstraZeneca)^5^. We observed no decrease in neutralisation titre with 501Y.V1 versus G614, compared to other leading vaccines, however host-specific effects may play a role^5^. It is encouraging that INO-4800-vaccinated ferret serum maintains neutralisation efficacy against the spectrum of emerging ‘variants-of-concern’ evaluated by this study^18^; this is consistent with the maintenance of neutralisation of pseudovirus prototypes of SARS-CoV-2 variants with serum samples from human recipients of the INO-4800 vaccine^19^. Our study indicates that nucleic acid-based vaccines will have an important role to play in this pandemic, in terms of their general ability to maintain neutralisation efficacy and to be modified for ‘vaccine matching’ in the future.

## Data Availability Statement

The data that support the findings of this study are available through the corresponding author upon reasonable request. Next generation sequencing data for the virus stocks is available from the NCBI BioProject database (BioProject ID: PRJNA722318).

## Acknowledgements

We are grateful for support from our colleagues at the Australian Centre for Disease Preparedness (https://www.grid.ac/institutes/grid.413322.5), particularly those in the Dangerous Pathogens and Animal Studies Teams who performed the parental vaccine study, and Kristen McAuley who facilitated importation of SARS-CoV-2 variants. The authors would like to thank the Victorian Infectious Diseases Reference Laboratory (especially Dr Mike Catton) for providing the VIC31 and VIC17990 (501Y.V1) isolates, and Tulio de Oliveira and Alex Sigal for providing the 501Y.V2.HV001 isolate. This work was conducted with funding to the Commonwealth Scientific and Industrial Research Organisation from the Australian Department of Finance, the Coalition for Epidemic Preparedness Innovations, and Inovio Pharmaceuticals, Inc.

## Author Contributions

A.J.M., P.A.D., and S.S.V. designed the study. S.R., S.G., A.J.M., and K.B. performed the experiments. A.J.M., P.A.D., and M.T. analysed the experimental data. M.J.K. performed biomolecular modelling and provided structural interpretations. J.D.D. isolated VIC31 and VIC17990 virus isolates; T.R.F.S. and K.E.B. co-invented the vaccine used in this study. A.J.M., P.A.D., M.J.K. and S.S.V. co-wrote the manuscript and made revisions. S.S.V. conceived the project and obtained funding. All authors reviewed the manuscript.

## Supplementary Figures

**Supplementary Figure 1.**
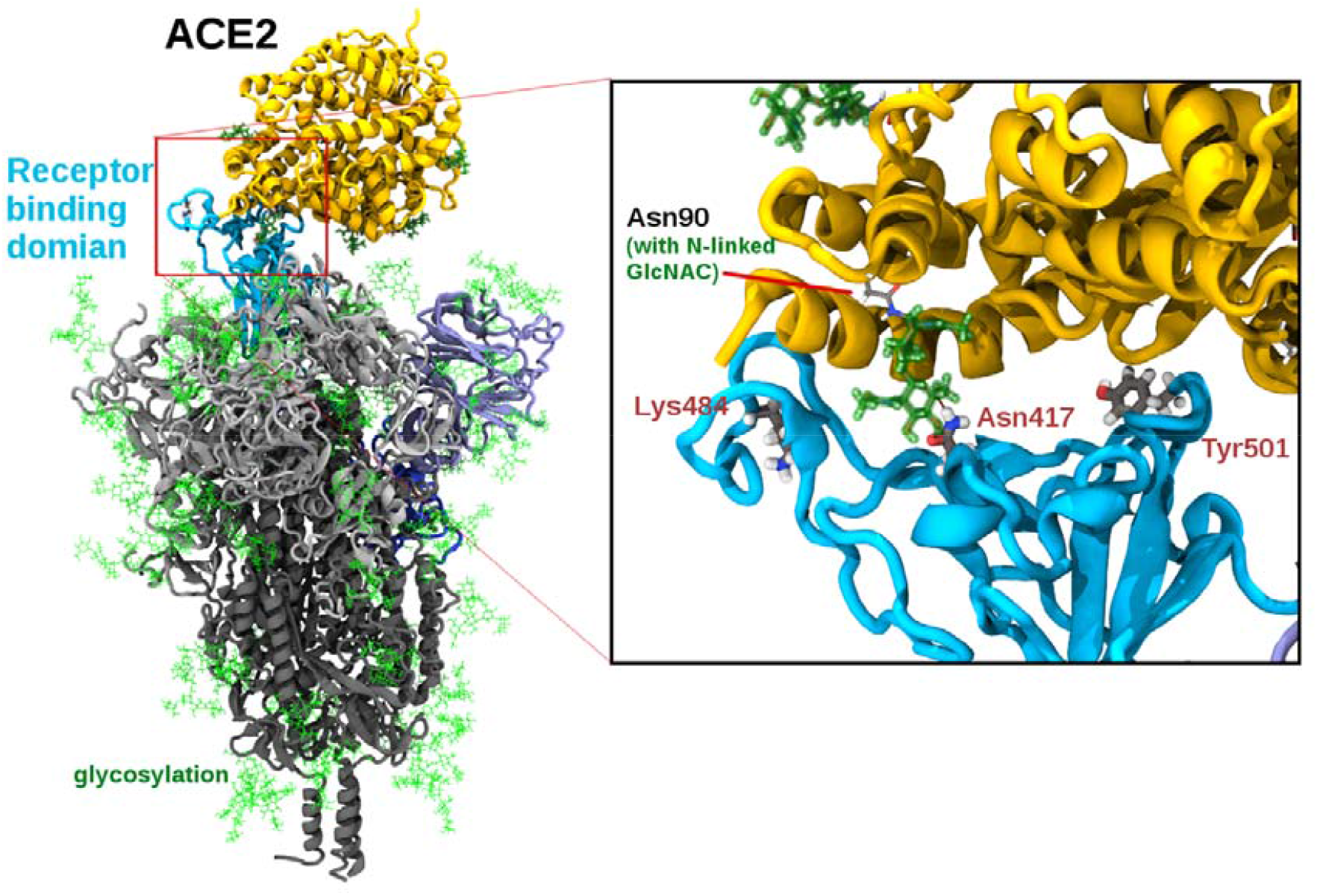
Structure of ACE2 binding to the SARS-CoV-2 Spike receptor-binding domain of variant 501Y.V2. Modelling suggests the lysine to asparagine mutation, (K417N), may facilitate additional hydrogen bonding to the ACE2 receptor, (Angeotensin-converting enzyme 2), via binding through an N-linked glycoside at position asparagine Asn90 in ACE2. (In this case modelled as Beta-D-GlcNAC (1->4) GlcNAC). The relative position of the other 501Y.V2 mutations in the receptor-binding domain (Tyr501 and Lys484) as also shown.

**Supplementary Figure 2.**
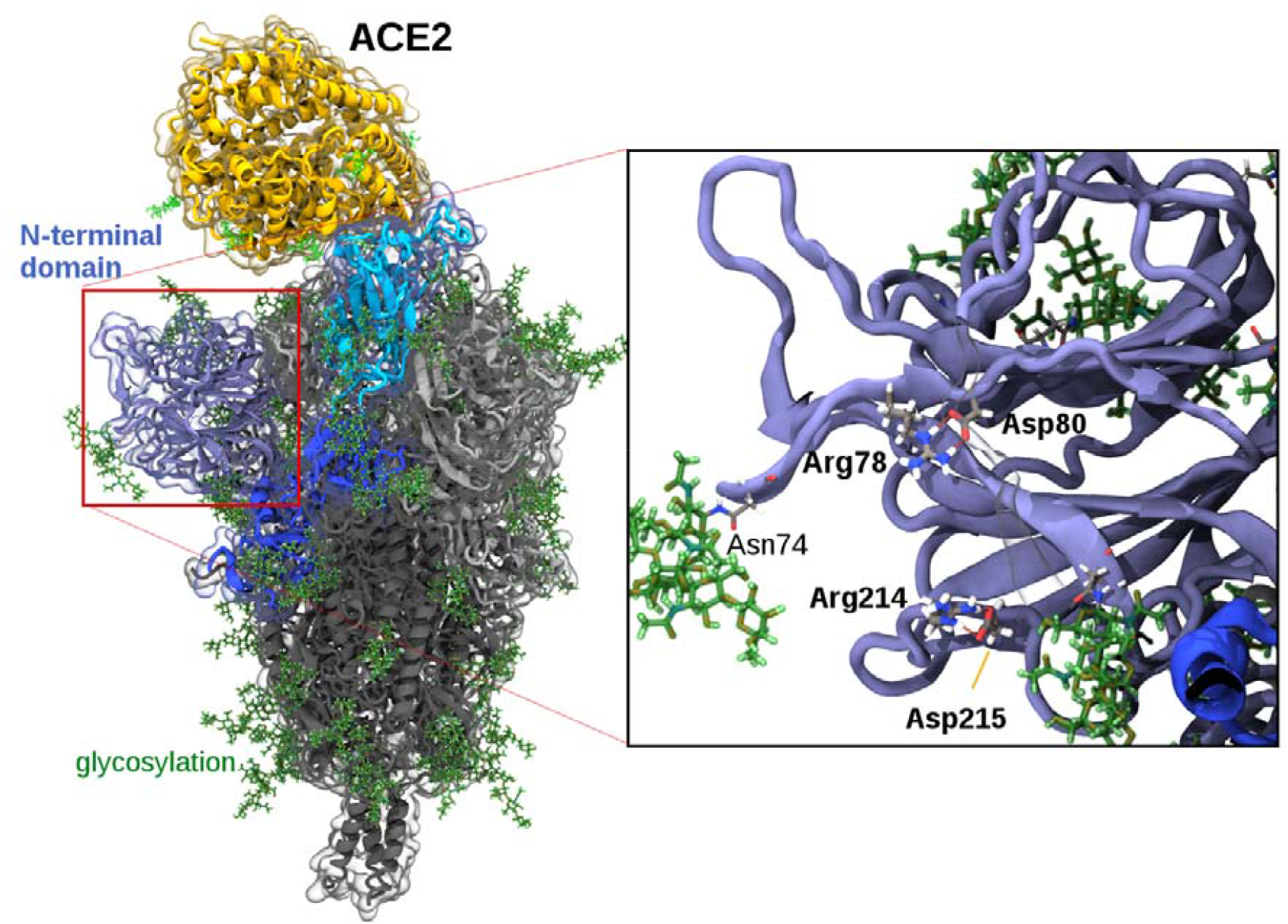
Structure of ACE2 binding to the SARS-CoV-2 Spike receptor-binding domain showing a close up of the N-terminal domain region, highlighting the salt bridges between arginine and aspartic acid at positions Arg78-Asp80 and Arg214-Asp215. In the 501Y.V2 variant, aspartic acid 80 is mutated to alanine (D80A) and aspartic acid 215 is mutated to glycine (D215G), losing both salt bridges, altering the relative positions of the arginine residues, as shown in Figure 2e). The asparagine N-linked glycosylation attachment point Asn74 is also labelled in the diagram.

